# Hypoxia disrupts circadian rhythms in astrocytes and causes synapse engulfment defects

**DOI:** 10.1101/2024.02.22.581651

**Authors:** Li Li, Jong Bin Choi, Chi Hong Shin, Saw Htun, Sherry Mestan, Anna Voss, Jennifer L. Shadrach, Alyssa Puno, Dhriti Nagar, Nephy Ramirez, Daniela Rojo, Samuel H. Lee, Erin M. Gibson, Julia A. Kaltschmidt, Steven A. Sloan, Won-Suk Chung, Anca M. Pasca

**Affiliations:** Department of Pediatrics, Stanford University, Palo Alto, CA 94305, USA; Department of Human Biology, Stanford University, Palo Alto, CA 94305, USA; Department of Biological Sciences, Korea Advanced Institute of Science and Technology (KAIST), Daejeon, Republic of Korea; Department of Human Genetics, Emory University, Atlanta, GA 30322, USA; Department of Neurosurgery, Stanford University, Palo Alto, CA 94305, USA; Department of Psychiatry and Behavioral Sciences, Stanford University, Palo Alto, CA 94305, USA; Department of Molecular and Cell Biology, University of California, Berkeley, Berkeley, CA, 94720

## Abstract

Astrocytes are emerging as key regulators of neuronal synaptic network maturation and function, through control of synaptic pruning. This is important, because individuals with ASD have excess glutamatergic synapses in the cortex, but the biological mechanisms underlying this phenotype remain unclear.

Here, we used human cortical organoids (hCO) derived from induced pluripotent stem cells (hiPSCs), to examine the effect of hypoxia on synapse engulfment in human astrocytes at postnatal-equivalent stages of development. We identified that hypoxia significantly inhibits the synaptosome phagocytosis, and that this phenotype is mediated through disruptions in the astrocytic circadian rhythm molecular pathway and subsequent decreased expression of MEGF10. Lastly, we demonstrated that circadian clock disruptions are sufficient to induce these observed phenotypes even in the absence of hypoxia, both in hCOs and within the mouse hippocampus *in vivo*.

Our study uncovers a novel mechanistic link between hypoxia, circadian rhythms disruptions, and synapse pruning by astrocytes, and provides insight into the pathophysiology of ASD, and other neuropsychiatric diseases. Separately, the demonstration of the presence of circadian rhythms in hCOs opens an unprecedented opportunity to dissect the role of circadian clocks in normal brain development and how it contributes to specific diseases of environmental or genetic origin.

## INTRODUCTION

Histological studies on postmortem brain tissue from individuals with autism spectrum disorders (ASD) have identified increased density of excitatory synapses in the cortex^1–3^. These findings align with functional studies (e.g. electroencephalography, functional MRI) demonstrating cortical hyperexcitability. One of the proposed hypotheses is a defect in physiologic synaptic pruning during brain development.

Traditionally, synaptic pruning had been known to be performed by microglia, which seed the developing brain during the first trimester of in utero development and represent about 5% of all cortical glia^4^. More recent studies have shown that astrocytes also actively phagocytose synapses and refine neural circuits in both developing and adult brain^5–8^.

Cortical astrocytes start developing in the second trimester of in utero development; their numbers progressively increase and eventually stabilize at about 30-50% of all cortical glia^9^. Interestingly, synapse pruning increases in parallel with the increasing number of astrocytes. These data suggest a possible under-appreciated role of astrocytes in physiologic synaptic pruning and underscore the need to investigate whether environmental insults clinically associated with increased risk for ASD (e.g. hypoxia and inflammation)^10–14^ affect this process.

Here, we investigated the effects of hypoxia exposure on the ability of human astrocytes to engulf synaptosomes. For this, we used astrocytes from human cortical organoids (hCOs) cultured *in vitro* for 10 months. At this time point hCO-derived astrocytes display postnatal characteristics^15,16^, a period in development when synaptic pruning is becoming increasingly more active.

For induction of hypoxia, we used our previously published method to study hypoxic brain injury of prematurity. Briefly, we exposed hCOs to partial pressure of oxygen ∼25-30 mmHg for 48 hr, which was enough to stabilize HIF1α and activate the hypoxia response molecular pathway^17^. We found that hypoxia exposure significantly decreases the ability of human astrocytes to engulf synaptosomes through a disruption of the cellular circadian clock and subsequent decreased expression of MEGF10, a key synaptic engulfment receptor in astrocytes^18^. Lastly, we validated the important role of circadian clocks in synaptic pruning by astrocytes using an *in vivo* rodent model.

Our study demonstrates for the first time the presence of circadian rhythms in hCOs and uncovers a novel mechanistic link between hypoxia, circadian clock disruption and synapse pruning defects in astrocytes. The identification of circadian clocks disruption in astrocytes as a risk factor for abnormal synaptic engulfment brings essential insight into pathophysiology of ASD and other developmental neuropsychiatric disorders and offers the opportunity to study circadian rhythm-modifier medications as possible therapeutic targets.

## RESULTS

### Hypoxia-exposed human astrocytes display synaptosome engulfment defects

To assess the effects of hypoxic stress on human astrocytes at early postnatal-equivalent stages of development, we used hCOs at 10 months in culture. In hCO-derived astrocytes, this time-point has been previously demonstrated to display postnatal transcriptional signatures when compared to human primary astrocytes^15,16^.

First, we confirmed the progression of astrocyte maturation in our set of experiments using hCOs generated from 4 individual human induced pluripotent stem cell lines (hiPSCs) and cultured for 6 and 10 months *in vitro*. At these time-points, we dissociated ∼40 hCOs/hiPSC line, isolated astrocytes by immunopanning using HepaCAM antibody^15,19^ and performed bulk RNA-sequencing (Extended Figure 1a). In line with previous reports using long-term culture hCOs^15^, we found that compared to 6-month astrocytes, astrocytes at 10-months upregulate the expression of genes enriched in mature astrocytes and downregulate those enriched in progenitor cells (Extended Figure 1b). In addition, we confirmed that astrocytes isolated from 10-month hCOs display more complex morphology, as shown by the increase in primary branch numbers when compared to astrocytes from 6-month hCOs (Extended Figure 1c and d). These data demonstrate that astrocytes generated for this study follow similar maturation patterns as previously shown, and are a valid in vitro model to study the effects of early-postnatal insults (e.g hypoxic ischemic encephalopathy) on astrocytes.

**FIGURE 1.**
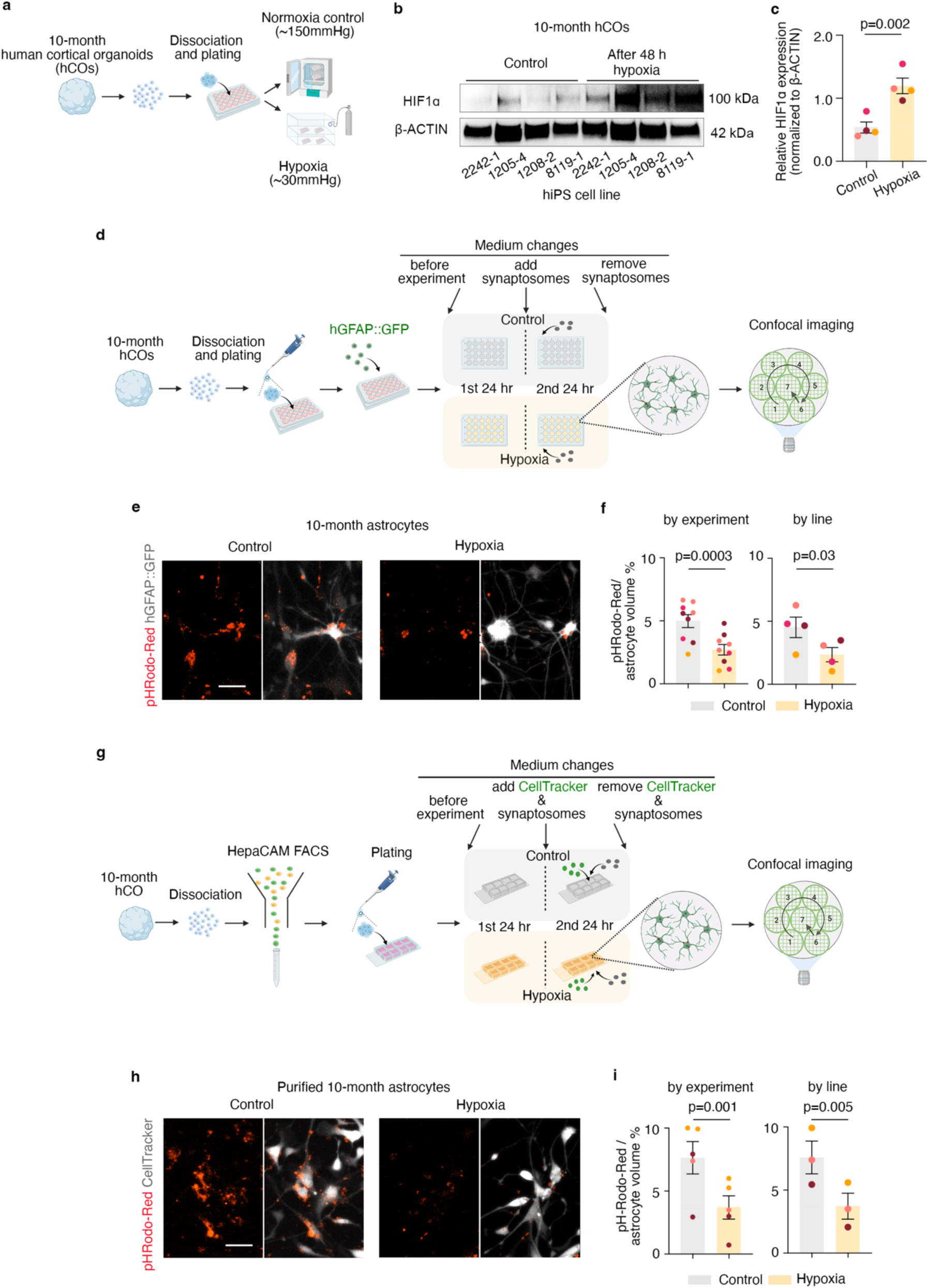
Hypoxia-exposed human astrocytes display synaptosome engulfment defects. **(a)** Schematic of differentiation of hCOs from hiPSCs, dissociation of 10-months hCOs and exposure to hypoxia; **(b)** Representative Western blot images of HIF1α and β-actin proteins in monolayer cultures of cells dissociated from 10-months hCOs, exposed to control or hypoxia condition for 48 hr; **(c)** Quantification of HIF1α in control and hypoxic conditions (two-tailed paired t-test; P=0.002); each color of dots represents one cell line; data are presented as mean ± s.e.m.; **(d)** Schematic showing synaptosome engulfment assay of astrocytes exposed to control or hypoxia condition; **(e)** Representative images of pHrodo-Red labeled synaptosomes engulfed (red) into 10-months astrocytes labeled with hGFAP::GFP lentivirus (gray) under control or hypoxia conditions; **(f)** Volume of synaptosomes engulfed by astrocytes under control or hypoxia conditions (normalized to total GFAP volume/well) analyzed by experiment (two-tailed paired *t*-test, P=0.0003) and by hiPSC line (two-tailed paired *t*-test, P=0.03); each condition was analyzed in 1-3 different differentiation experiments; **(g)** Schematic showing astrocyte purification by FACS using HepaCAM antibody followed by synaptosome engulfment assay of enriched astrocytes exposed to control or hypoxia conditions; **(h)** Representative images of pHrodo-Red labeled synaptosomes engulfed (red) into 10-months astrocytes purified by FACS using HepaCAM antibody and labeled with CellTracker (gray) under control or hypoxia conditions; **(i)** Volume of synaptosomes engulfed by 10-months astrocyte purified by HepaCAM FACS (normalized to total CellTracker volume) analyzed by experiment (two-tailed paired *t*-test, P= 0.001) and by hiPSC line (two-tailed paired *t*-test; P=0.005); each experiment was performed on astrocytes purified from ∼15 hCOs/hiPSC line, in 1-2 experiments, using 3 hiPSC lines. All the hiPSC lines and hCO used in experiments and figures are summarized in Extended Table 1. Each color of dots represents one cell line. Data are presented as mean ± s.e.m.

To establish a model for hypoxic astrocyte injury, we dissociated 10-months hCOs, plated them in monolayer and cultured them for ∼3-5 days. Subsequently, we exposed the cells to control conditions (∼150 mmHg, 5% CO2, 37 °C) or to hypoxia (∼30 mmHg, 5% CO2, 37 °C) for a total of 48 hr (Figure 1a). Using an optical microsensor of oxygen tension (OXB50, PyroScience) attached to a fiber-optic multi-analyte meter (FireStingO_2_, PyroScience), we measured the partial pressure of oxygen (P*O_2_*) in the control and hypoxic medium, and validated the expected decrease of P*O_2_* (Extended Figure 1e). We confirmed that these conditions are sufficient to induce stabilization of HIF-1α (two-tailed paired *t*-test, P=0.002) (Figure 1b and c), as expected based on our previously published data^17,20^.

Next, we examined whether hypoxia exposure affects the capability of hCO-derived astrocytes to engulf synaptosomes *in vitro*. To do this, we dissociated ∼12 hCOs/hiPSC line, plated them as a dot-shape monolayer at the center of the well in 24 well plates, and cultured them for ∼3-5 days. To label astrocytes in the monolayer culture, we infected the cells with hGFAP::GFP lentivirus, where GFP expression is driven by the human GFAP promoter. After another 3 days we observed robust expression of GFP within cells with astrocytic morphology. At this time-point, we exposed cells to control conditions (∼150 mmHg, 5% CO2, 37 °C) or to hypoxia (∼30 mmHg, 5% CO2, 37 °C) for a total of 48 hr. After 24 hr, we supplied fresh culture medium supplemented with pHrodo-Red-labeled synaptosomes to both control- and hypoxia-exposed cells. At the end of 48 hr treatment, we performed the last medium change to remove synaptosomes, and proceeded immediately with confocal microscopy imaging. Images were taken counterclockwise to first cover the cells at the edge then the center of the dot-shape area for both control- and hypoxia-exposed cells (Figure 1d).

We identified that hypoxic exposure significantly reduces the amount of synaptosomes engulfed by astrocytes (by experiment: two-tailed paired *t*-test, P=0.0003; by hiPSC line: two-tailed paired *t*-test, P=0.03) (Figure 1e and f). For these analyses, we used Imaris to quantify the volume of synaptosomes engulfed (pHrodo-Red^+^) into GFP-tagged cells in each field being imaged. To exclude the possible variation caused by cell density, we normalized pHrodo-Red to the total volume of GFP^+^ cells in each field.

To confirm that the synaptosomes showing fluorescence are indeed phagocytosed into astrocytes, we performed immunostaining for synaptosomes (mCherry) and the lysosome marker LAMP2 within hGFAP::GFP labeled astrocytes (GFP^+^). As expected, we found strong colocalization of synaptosomes with LAMP2 in both control and hypoxia conditions (Extended Figure 1g).

To validate this phenotype using a pure population of astrocytes, we first enriched astrocytes from 10-month hCOs by FACS using HepaCAM antibody, as described previously^15^. We cultured the enriched astrocytes in monolayer for 3 days, and then exposed them to control or hypoxia conditions for a total of 48 hr (Figure 1g). For imaging, we labeled the enriched astrocytes using CellTracker CMFDA and supplemented pHrodo-Red labeled synaptosomes in the last 24hr of hypoxia treatment, as described in the previous experiments. Consistent with the results in dissociated hCOs, we observed significant reduction of synaptosome engulfment by astrocytes upon exposure to hypoxia (by experiment: two-tailed paired *t*-test, P=0.001, by hiPSC line: two-tailed paired *t*-test, P=0.005) (Figure 1h and i).

These results demonstrate that hypoxia exposure substantially decreases the capability of human astrocytes to engulf synaptosomes.

### Circadian rhythm pathway-associated genes are dysregulated in hypoxic human astrocytes

To investigate hypoxia-induced transcriptional changes in hCO-derived astrocytes, we first validated the induction of hypoxia in intact hCOs exposed to ∼30 mmHg (5% CO2, 37 °C) for 48 hr, by using HIF-1α protein stabilization (two-tailed paired *t*-test, P=0.001) (Figure 2a and b). These data align with our previous reports using hCOs at early stages of differentiation^17,20^.

**FIGURE 2.**
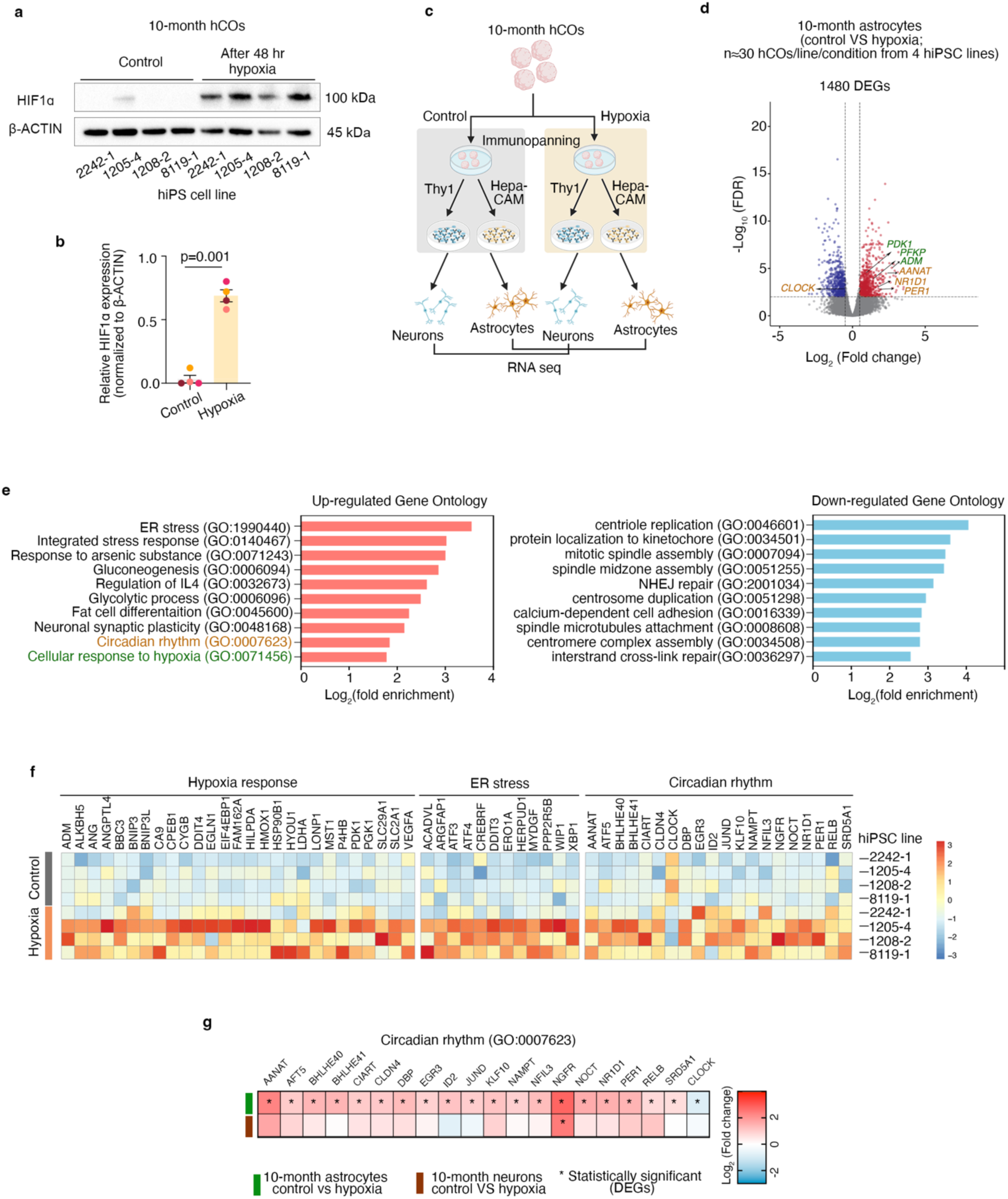
Circadian clock genes are dysregulated in human astrocytes under hypoxic stress. **(a)** Representative Western blot images of HIF1α and β-actin proteins in intact 10-month hCOs exposed to control or hypoxia condition for 48 hr; **(b)** Quantification of HIF1α in control and hypoxic conditions (two-tailed paired *t*-test; P=0.001); **(c)** Schematic of experimental design for exposure to hypoxia of intact hCOs, and purification by immunopanning of astrocytes and neurons used for RNA-sequencing; **(d)** Volcano plots of differentially expressed genes in RNA-seq of astrocytes after exposure to hypoxia for 48 hr compared to control; each dot represents a single gene, with significantly upregulated genes shown in red, significantly downregulated in blue, and non-significant in gray (FDR<0.01 and |Log_2_fold change| ≥ 0.5 as cutoff); n=40-50 hCOs/hiPSC line, total 4 hiPSC lines for each time point. Complete list of all DEGs can be found in Extended Table 2; **(e)** Bar plots of the top upregulated biological processes identified by Gene ontology (GO) analysis of DEGs associated with hypoxia; complete list of all GO terms can be found in Extended Table 3. **(f)** Heatmaps of TPM values of DEGs identified in corresponding GO terms; **(g)** Heatmaps of Log_2_fold changes of DEGs identified within the circadian rhythm pathway in astrocytes (green bar to the left of the map) and neurons (brown bar to the left of the map); stars(*) on the heatmap indicate statistically significant in the comparison between control and hypoxia (FDR<0.01 and |Log_2_fold change| ≥ 0.5 as cutoff). A detailed list of FDR (p-adj) values and Log_2_fold changes for all the listed genes can be found in Extended Tables 4 and 5.

Using these experimental conditions, we performed bulk RNA-seq on astrocytes enriched by immunopanning using HepaCAM antibody from 10-months hCOs derived from 4 hiPSC lines (Figure 2c), exposed and non-exposed to hypoxia. We identified 1480 DEGs (794 up-regulated and 686 down-regulated) (FDR<0.01 and |Log_2_fold change| ≥ 0.5) (Figure 2d, Extended Table 2). Top biological processes affected by hypoxia were identified using Gene Ontology (GO) analysis. Intriguingly, circadian rhythm (GO:0007623) was one of the top GO terms (Figure 2e and f). While it has been previously reported in skeletal muscle cells and tumor cells that hypoxia induces circadian rhythm pathway disruptions through HIF1α^21–23^, this regulatory connection has not been yet identified in astrocytes. As expected, we also identified transcriptional induction of cellular response to hypoxia (GO: 0071456), glycolysis (GO: 0061621 and GO: 0006096), gluconeogenesis (GO: 0006094) and endoplasmic reticulum (ER) stress pathway (GO:0070059 and GO:1990440)^4^ and downregulation of genes associated with cell replication (Figure 2e and f; Extended Tables 3 and 4).

To check if circadian pathway disruption is cell type-specific, we analyzed neurons immunopanned from the same 10-months hCOs using THY1 antibody^15,19^. Interestingly, we found that neurons show minimal changes in the expression of circadian rhythm-related genes (Figure 2g; Extended Tables 4 and 5).

Taken together, these results uncover a significant and cell-type specific response of circadian rhythm-related genes to hypoxia in astrocytes.

Due to our protocol setup, all the processes of immunopanning were performed around a similar time of the experiment day. Specifically, hCOs under both control and hypoxia conditions were dissociated in the mornings around 8:30-9:00 AM. Immunopanning experiments were completed and cells were snap-frozen for RNA isolation around 3:00-4:00 PM of the day. The synaptosome engulfment assays in Figure 1 were also performed in the early afternoon around 2:00-4:00 PM. As the timing of sample collection and performing the assays are critical for the assays involving circadian rhythm, all the follow-up experiments were performed at these times.

### Circadian rhythms are functionally disrupted in hCOs exposed to hypoxia

Circadian rhythms are regulated by a transcriptional-translational-feedback loop (TTFL) consisting of key clock genes, such as *CLOCK*, *BMAL*, *PER*s and CRYs^24^. REV-ERBα (encoded by *NR1D1)* is a key component of the secondary regulatory loop of the molecular circadian clock through inhibiting transcription of *BMAL1*^25,26^(Figure 3a).

**FIGURE 3.**
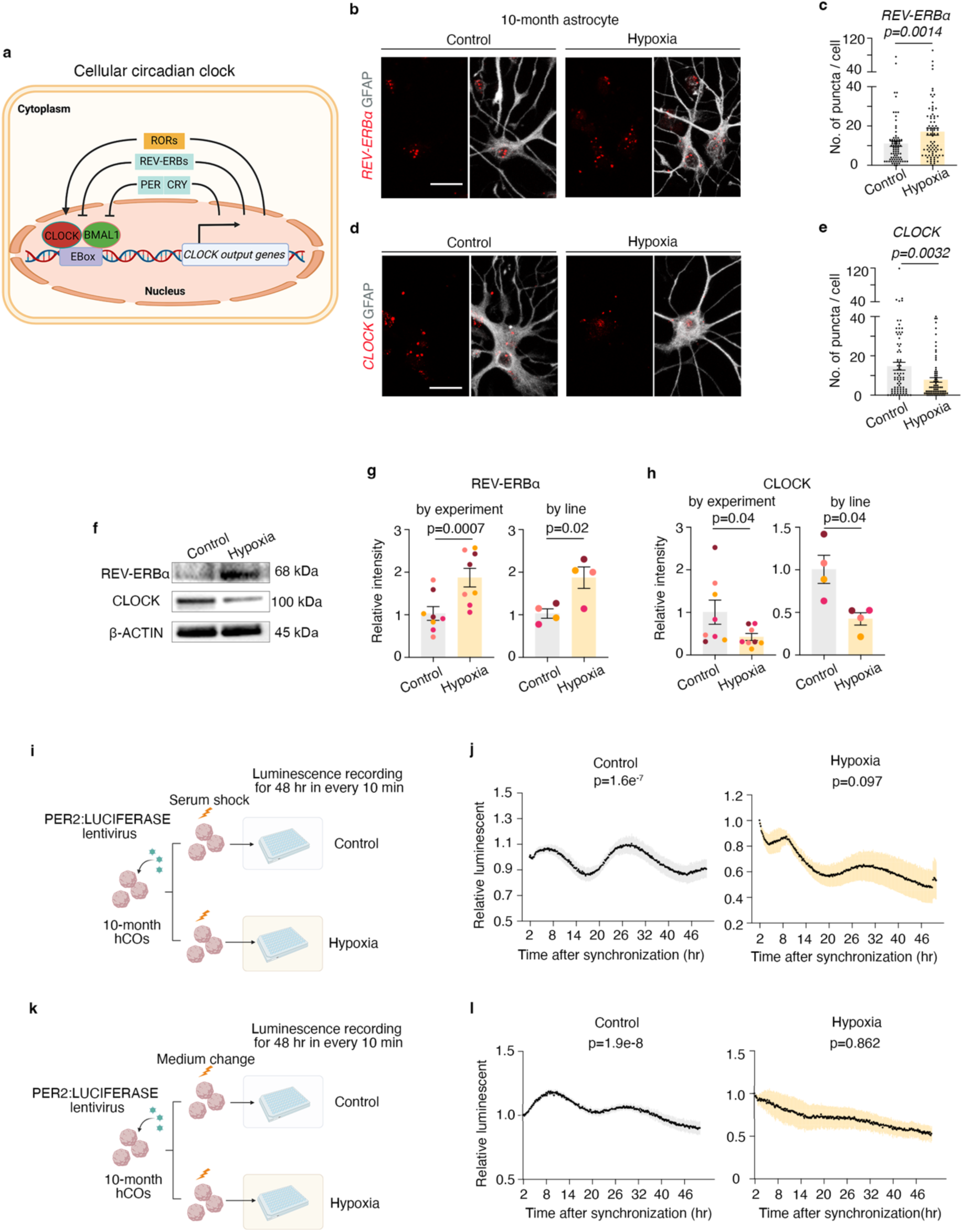
Circadian rhythm is disrupted in hCOs and astrocytes exposed to hypoxia. **(a)**Schematic of circadian clock machinery; **(b**) Representative images of in situ RNA hybridization of *REV-ERBα* (red) within GFAP^+^ astrocytes (gray) in control and hypoxia; **(c)** Quantification of the numbers of in situ *REV-ERBα* transcripts per GFAP^+^ astrocytes after exposure to control or hypoxia (unpaired Mann-Whitney test, P=0.0014); n=25-80 cells isolated from ∼30 hCOs/hiPSC line, total 2 hiPSC lines; each dot represents one GFAP^+^ cell quantified for transcript number; **(d)** Representative images of in situ RNA hybridization of *CLOCK* (red) and GFAP^+^ astrocytes (gray) in control and hypoxia; **(e)** Quantification of the numbers of in situ *CLOCK* transcripts per GFAP^+^ astrocytes after exposure to control or hypoxia (unpaired Mann-Whitney test, P=0.0032); n=25-80 cells isolated from ∼30 hCOs/hiPSC line, total 2 hiPSC lines; each dot represents one GFAP^+^ cell quantified for transcript number; scale bar: 20 μm; **(f)** Representative Western blot images of REV-ERBα, CLOCK and β-actin proteins in 10-months hCOs exposed to control or hypoxia; **(g)** Quantification of REV-ERBα in control and hypoxic conditions, analyzed by experiment (two-tailed paired *t*-test; P=0.0007) and by hiPSC line (two-tailed paired *t*-test; P=0.02); each color of dots represents one hiPSC line; data are presented as mean ± s.e.m.; **(h)** Quantification of CLOCK in control and hypoxic conditions, analyzed by experiment (two-tailed paired *t*-test; P=0.04) and by hiPSC line (two-tailed paired *t*-test; P=0.04); each color of dots represents one cell line; data are presented as mean ± s.e.m.; **(i)** Schematic showing PER2::LUCIFERASE lentiviral infection and assay in intact hCOs synchronized by serum shock; **(j)** 48 hr luminescence recording of intact 10-months hCOs infected with PER2::LUCIFERASE, after synchronization with serum shock, and exposed or non-exposed to hypoxia (n=3-9 hCOs/hiPSC line in control condition, from 3 hiPSC lines derived by 3 differentiations; n=6 hCOs/hiPSC line in control condition, from 2 hiPSC lines derived by 2 differentiations). Relative luminescence intensity was calculated by normalizing to the 13th time point (T13, 2 hr after recording started). Rhythmicity was analyzed using JTK_cycle analysis (ANOVA control P=1.6e-7; hypoxia P=0.097). **(k)** Schematic showing PER2::LUCIFERASE lentiviral infection and assay in intact hCOs synchronized by medium change; **(l)** 48 hr luminescence recording every 10 min of intact 10-months hCOs infected with PER2::LUCIFERASE, after synchronization with medium change, and exposed or non-exposed to hypoxia (n=3 hCOs/hiPSC line in each condition, total 2 hiPSC lines). Relative luminescence intensity was calculated by normalizing to the 13th time point (T13, 2 hr after recording started). Rhythmicity was analyzed using JTK_cycle analysis (ANOVA control P=1.9e-8; hypoxia P=0.862). Data are presented as mean (black line) and error bar represents s.e.m. (yellow).

To validate the transcriptional changes of circadian genes observed in RNA-seq in hypoxic astrocytes, we used multiple assays. First, we used RNAscope in astrocytes derived from 10-months hCOs and cultured in monolayer, to check the transcript levels of *CLOCK -* a central driver of circadian rhythm, and *REV-ERBα* -a negative regulator of the of the circadian clock. Consistent with RNA-seq results, we observed a significant increase of *REV-ERBα* transcript level under hypoxic conditions (unpaired Mann-Whitney test, P=0.0014) (Figure 3b and c). On the other hand, *CLOCK* was significantly down-regulated in hypoxic astrocytes (unpaired Mann-Whitney test, P=0.0032) (Figure 3d and e). To check whether the transcriptional changes translate into protein level alterations, we performed Western blot analyses for both REV-ERBα and CLOCK in 10-months hCOs. In line with the transcriptional changes, we identified increased expression of REV-ERBα when analyzed by experiment (two-tailed paired *t*-test, P=0.0007) and by hiPSC line (two-tailed paired *t*-test, P=0.02) ) (Figure 3f and g), and decreased protein expression of CLOCK when analyzed by experiment (two-tailed paired *t*-test, P=0.04) and by hiPSC line (two-tailed paired *t*-test, P=0.04) (Figure 3f and h).

To investigate whether these gene/protein specific changes correlate with functional alterations in intact hCOs, we used PER2::LUCIFERASE lentivirus, which is commonly used to detect circadian rhythm in animal models, primary cells or immortalized cell lines^8–10^. One week after viral infection of 10-months hCOs, we used serum shock to synchronize the cells for 2 hr followed by luminescence recording every 10 min under control or hypoxia conditions for another 48 hr (Figure 3i). Time points from the 13^th^ recording (T13, 2 hr after recording started) were included for analyses as luminescence signal gets stabilized from this time point. Luminescence intensity at each time point from both control and hypoxia conditions was normalized to beginning time point (T13). Normalized time series were analyzed by JTK analysis using NiteCap, an online platform for quantifying circadian and rhythmic behavior^27^. Using serum shock, we observed strong rhythmicity in hCOs under control conditions (ANOVA P=1.6e-7). Upon hypoxia exposure (∼25-30mmHg, Extended Figure 3a), hCOs lost this rhythmicity (ANOVA P=0.097) (Figure 3j). As we performed media changes for synaptosome engulfment assays, we next tested whether medium change was sufficient to synchronize the cells. With medium change before luminescence recording, we also observed robust rhythmicity in hCOs ( ANOVA P=1.9e-8). This rhythmicity is dramatically disrupted upon hypoxia exposure (ANOVA P=0.862) (Figure 3k and l).

These results identify, for the first time, the presence of circadian rhythms in hCOs, and uncover the disruption of this important process by exposure to hypoxia.

### Downregulation of *REV-ERBα* is sufficient to rescue the synaptosome engulfment defect in hypoxic astrocytes

Given that disruptions in circadian rhythms are strongly associated with ASD in the clinical setting, and histological samples from the brain of individuals with ASD demonstrate an excess of excitatory synapses, we went on to further investigate whether circadian rhythm manipulations in astrocytes can affect synaptosome engulfment in the presence or absence of hypoxia.

*REV-ERBα (NR1D1*) is a key repressor of circadian clock and gatekeeps a wide range of cellular functional outputs of circadian clock^26,28^. In addition, *REV-ERBα* was up-regulated dramatically in hypoxic astrocytes. We decided to investigate whether decrease in REV-ERBα level is sufficient to rescue the synaptosome engulfment defect in hypoxic astrocytes. To address this question, we performed siRNA knockdown of *REV-ERBα* using 2 siRNAs targeting two different sequences at Exon 1 of *REV-ERBα* (si*REV-ERBα*-1 and si*REV-ERBα*-2) in hypoxic astrocytes (Figure 4a). We first demonstrated a reduction in *REV-ERBα* mRNA level by both siRNAs treatment (one-way ANOVA; si*REV-ERBα*-1: P=0.035; si*REV-ERBα*-2: P=0.014) compared to siRNA scramble control (siScramble) (Extended Figure 4a), and validated the protein level decrease in HEK 293T cells (Extended Figure 4b and c). Next, we transfected monolayer cultures of enriched 10-month hCO-derived astrocytes with siScramble, si*REV-ERBα*-1 and si*REV-ERBα*-2 and confirmed the decrease in *REV-ERBα* expression in astrocytes (one-way ANOVA; si*REV-ERBα*-1: P=0.033; si*REV-ERBα*-2, P=0.014) (Extended Figure 4d). Using these validated siRNAs, we performed synaptosome engulfment assays under hypoxia condition according to the protocol described previously. We identified significant rescue of synaptosome engulfment in hypoxic astrocytes with both si*REV-ERBα*-1 and -2 compared to siScramble, when analyzed by experiment (one-way ANOVA; si*REV-ERBα*-1: P=0.0007; si*REV-ERBα*-2: P=0.0015) and by hiPCS line (one-way ANOVA; si*REV-ERBα*-1: P=0.012; si*REV-ERBα*-2: P=0.031) (Figure 4b and c).

**FIGURE 4.**
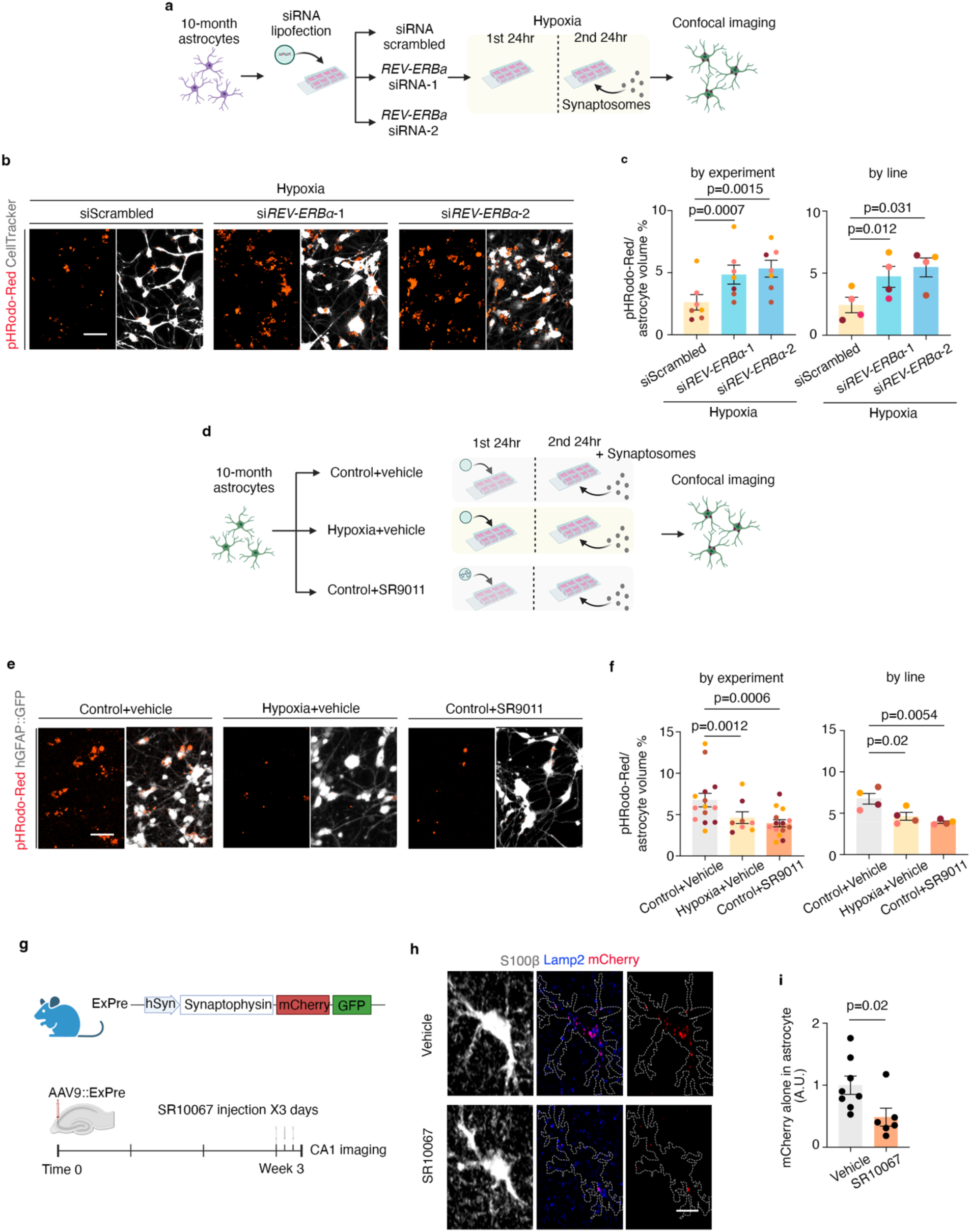
Regulation of synapse engulfment by REV-ERBα in human and mouse astrocytes. **(a)** Schematic of siRNA transfection protocol and synaptosome engulfment assay in purified astrocytes, following exposure to hypoxia; **(b)** Representative images of synaptosomes (red) within hypoxic astrocytes (gray) transfected with siRNA scramble control (siScramble), si*REV-ERBα*-1 or si*REV-ERBα*-2; scale bar: 20 μm; **(c)** Quantification of synaptosome engulfment by hypoxic purified astrocytes transfected with siScramble versus si*REV-ERBα*-1 and si*REV-ERBα*-2 analyzed by experiment (one-way ANOVA; si*REV-ERBα*-1: P=0.0007; si*REV-ERBα*-2: P=0.0015) and by hiPCS line (one-way ANOVA; si*REV-ERBα*-1:P=0.012; si*REV-ERBα*-2: P=0.031); **(d)** Schematic of experimental design for treating purified astrocytes with SR9011, a REV-ERBα agonist to compare synaptosome engulfment in control, hypoxia and SR9011-treated cells; **(e)** Representative images of synaptosomes (red) within hypoxic astrocytes (gray) in control treated with drug dissolvent (Control+Vehicle), hypoxia treated with drug dissolvent (Hypoxia+Vehicle) and Control+SR9011 conditions; scale bar: 20 μm; **(f)** Quantification of synaptosome engulfment by astrocytes under control condition compared to hypoxia condition (by experiment: mixed-effects analyses; P=0.0012; by hiPSC line: paired one-way ANOVA, P=0.02), and compared to Control+SR9011 treatment (by experiment: mixed-effects analyses; P=0.0006; by hiPSC line: paired one-way ANOVA, P=0.0054); **(g)** Schematic of in vivo adult mice treatment with SR10067, a REV-ERBα agonist, within the hippocampus; **(h)** Representative images of synapses engulfed (mCherry) into astrocytes (S100β staining; gray), co-stained with lysosome marker Lamp2 (blue) in Vehicle- and SR10067-treated mouse brains; scale bar: 10 μm; **(i)** Quantification of synapses engulfed into astrocytes (unpaired two-tailed t-test, P=0.02) (n= 4, 3 mice per condition, each dot represents 1 imaging field and two images were taken in each mouse brain). Each color of dots represents one cell line for hCO experiments. Data are presented as mean ± s.e.m.

These results suggest *REV-ERBα* manipulation is sufficient to restore the synapse engulfment defects in astrocytes caused by hypoxia exposure.

### Pharmacological increased biological activity of REV-ERBα is sufficient to decrease synaptosome intake by human astrocyte in the absence of hypoxia

To investigate whether disruption of circadian rhythm inhibits synaptosome engulfment by astrocytes independently from hypoxia, we treated astrocytes with SR9011 (10 μM)^29–31^, an agonist known to increase the biological activity of REV-ERBα, and performed synaptosome engulfment assay (Figure 4d). First, we checked the protein expression of REV-ERBα and CLOCK in the presence of SR9011. As expected, we did not see a change in REV-ERBα protein level (by experiment: paired Wilcoxon test, P=0.312; by hiPSC line: paired Wilcoxon test, P=0.375) (Extended Figure 4e and f); we did however identify decreased protein level of CLOCK, suggesting the dose of SR9011 used here was sufficient to disrupt the progression of circadian clocks in astrocytes (by experiment: two-tailed paired *t*-test, P<0.0001; by hiPSC line: two-tailed paired *t*-test, P=0.002) (Extended Figure 4e and g).

Next, we found that addition of SR9011 to control astrocytes significantly decreases the synaptosome engulfment compared to vehicle-treated condition (by experiment: mixed-effects analyses; P=0.0006; by hiPSC line: paired one-way ANOVA, P=0.0054), mimicking the effects of hypoxia (by experiment: mixed-effects analyses; P=0.0012; by hiPSC line: paired one-way ANOVA, P=0.02) (Figure 4e and f).

These results suggest that REV-ERBα, a key component of circadian clock machinery, directly regulates synaptosome engulfment by astrocytes.

### Regulation of synapse engulfment by REV-ERBα is conserved in adult mouse hippocampal astrocytes

Synapse turnover is critical for neurodevelopment as well as adult brain plasticity^5^. To investigate whether increase of REV-ERBα activity, with subsequent disrupted cellular circadian clock progression, also regulates synapse phagocytosis *in vivo* in the mouse brain, we injected AAV virus carrying hSYN promoter driven phagocytosis reporter AAV9-ExPre SYP-mCherry-eGFP to target the presynaptic structures of excitatory synapses in mouse hippocampus^5^. Excitatory synapses phagocytosed into lysosomes preserve the mCherry signal but not the eGFP signal thus allowing us to identify excitatory synapses engulfed into astrocytes.

The AAV reporter was injected into the CA3 region of the adult mouse hippocampus and these mice were treated with REV-ERB agonist SR10067^30^ or Vehicle control for 3 consecutive days (Figure 4g). Mouse brains were harvested and processed for analysis. We found *Clock* level in mouse brain was decreased following SR10067 injections compared to Vehicle treatment, suggesting repressing effects on circadian clock by SR10067 treatment as expected (two-tailed unpaired *t*-test, P<0.0001) (Extended Figure 4h). To examine the effect of SR10067 on synapse phagocytosis, we immunostained astrocytes with S100β and labeled lysosome with Lamp2 and engulfed synapses with mCherry. We observed mCherry-alone puncta co-localized with lysosome marker Lamp2 inside S100β^+^ astrocytes, demonstrating that they are indeed phagocytosed into astrocytes (Figure 4h). Next, we quantified fluorescence intensity of mCherry-alone puncta in S100β^+^astrocytes. Consistent with *in vitro* results, SR10067 administration significantly decreased the amount of synapse engulfed into mouse hippocampal astrocytes (two tailed Mann-Whitney test, P=0.02) (Figure 4h, i).

Taken together with *in vitro* hCO data, these results suggest REV-ERBα directly regulates synapse phagocytosis by astrocytes, and this regulation is conserved across human and mouse species and across ages.

### MEGF10 acts downstream of REV-ERBα to decrease synapse engulfment by human astrocytes

MEGF10 and MERTK are two classical phagocytosis receptors for astrocytes in rodent models^18^ (Figure 5a). From the RNA-seq comparison between control and hypoxic astrocytes, we found that *MEGF10*, but not *MERTK* mRNA level significantly decreased in astrocytes upon hypoxia exposure (*MEGF10*: two-tailed paired *t*-test, P=0.02; *MERTK* (two-tailed paired *t*-test, P=0.82) (Figure 5b). To further confirm this at protein level, we performed Western blot using 10-months hCOs exposed to control or hypoxia conditions for 48 hr. Consistent with RNA-seq results, MEGF10 protein level was decreased upon hypoxia treatment when analyzed by experiment (two-tailed paired *t*-test, P=0.0003) and by hiPCS line (two-tailed paired *t*-test, P<0.0001) (Figure 5c and d), while MERTK expression remained stable when analyzed by experiment (two-tailed Mann Whitney test, P=0.959) and by hiPCS line (two-tailed paired *t*-test, P=0.07) (Figure 5c and e).

**FIGURE 5.**
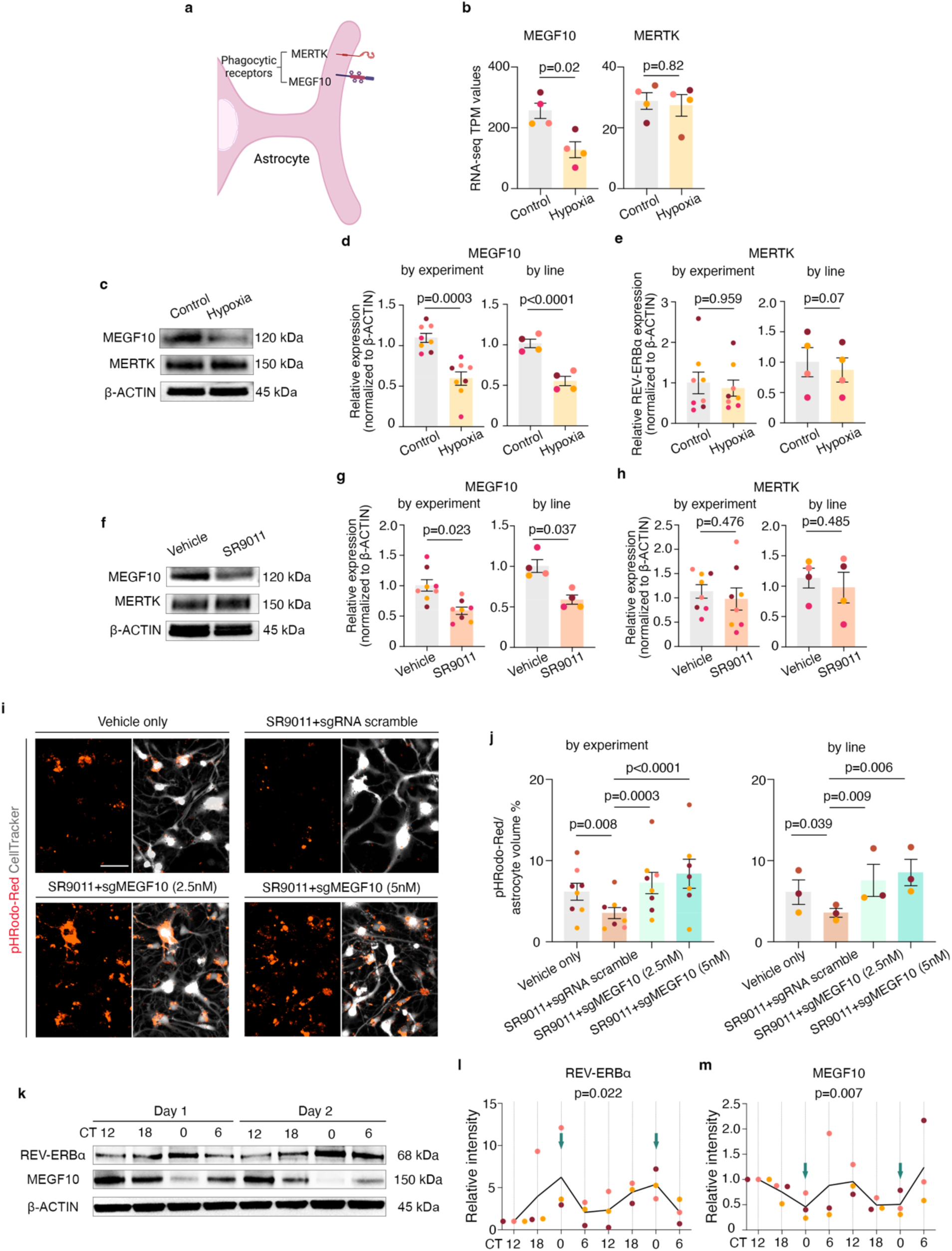
Regulation of synapse engulfment by REV-ERBα through MEGF10 in human astrocytes. **(a)** Schematic showing phagocytosis receptors MEGF10 and MERTK on astrocytes. **(b)** TPM values (from RNA-seq data) of MEGF10 (two-tailed paired *t*-test, P=0.02) and MERTK (two-tailed paired *t*-test, P=0.82) under control and hypoxia conditions expression; **(c)** Representative Western blot images of MEGF10, MERTK and β-actin proteins in 10-months hCOs exposed to control or hypoxia condition; **(d)** Quantification of MEGF10 protein level relative to β-ACTIN in control and hypoxic conditions, analyzed by experiment (two-tailed paired *t*-test, P=0.0003) and by hiPSC line (two-tailed paired *t*-test, P<0.0001); **(e)** Quantification of MERTK protein level relative to β-ACTIN in control and hypoxic conditions, analyzed by experiment (two-tailed Mann Whitney test, P=0.959) and by hiPSC line (two-tailed paired *t*-test, P=0.07); **(f)** Representative Western blot images of MEGF10, MERTK and β-actin proteins in 10-months hCOs treated with Vehicle or SR9011; **(g)** Quantification of MEGF10 protein levels relative to β-ACTIN in Vehicle or SR9011 treatment conditions, analyzed by experiment (two-tailed paired *t*-test; P=0.023) and by hiPSC line (two-tailed paired *t*-test; P=0.037); **(h)** Quantification of MERTK protein levels relative to β-ACTIN in Vehicle or SR9011 treatment conditions,, analyzed by experiment (two-tailed paired *t*-test; P=0.476) and by hiPSC line (two-tailed paired *t*-test; P=0.485); **(i)** Representative images of pHrodo-Red labeled synaptosomes engulfed (red) into 10-months astrocytes and labeled with CellTracker (gray) in vehicle ethanol treated cells (Vehicle only), SR9011-treated cells transfected with sgRNA non-targeting pool (SR9011+sgRNA scramble), SR9011-treated cells transfected with sgRNA pool targeting MEGF10 promoter at 2.5nM (SR9011+sgMEGF10 2.5nM), and SR9011treated cells transfected with sgRNA targeting MEGF10 promoter at 5nM (SR9011+sgMEGF10 5nM); **(j)** Volume of synaptosomes engulfed by 10-months astrocytes normalized to total CellTracker volume in Vehicle control versus SR9011+sgRNA scramble (by experiment: mixed-effects analysis, P=0.008; by hiPSC line: RM one-way ANOVA; P=0.039), SR9011+sgMEGF10 2.5M (by experiment: mixed-effects analysis, P=0.0003; by hiPSC line: RM one-way ANOVA; P=0.009), and SR9011+sgMEGF10 5nM (by experiment: mixed-effects analysis, P<0.0001; by hiPSC line:_RM one-way ANOVA; P=0.006); **(k)** Representative Western blot images of REV-ERBα, MEGF10 and β-actin proteins in hCOs synchronized by medium change and collected every 6 hr over a total of 48 hr; **(l)** Quantification of REV-ERBα protein level relative to β-ACTIN at corresponding time point over 48 hr (JTK_Cycle analysis, P=0.022); **(m)** Quantification of MEGF10 protein level relative to β-ACTIN at corresponding time point over 48 hr (JTK_Cycle analysis, P=0.007). Arrows (green) point to CT 0 time points (12AM) on each plot. Each color of dots represents one cell line; data are presented as mean ± s.e.m.

**FIGURE 6.**
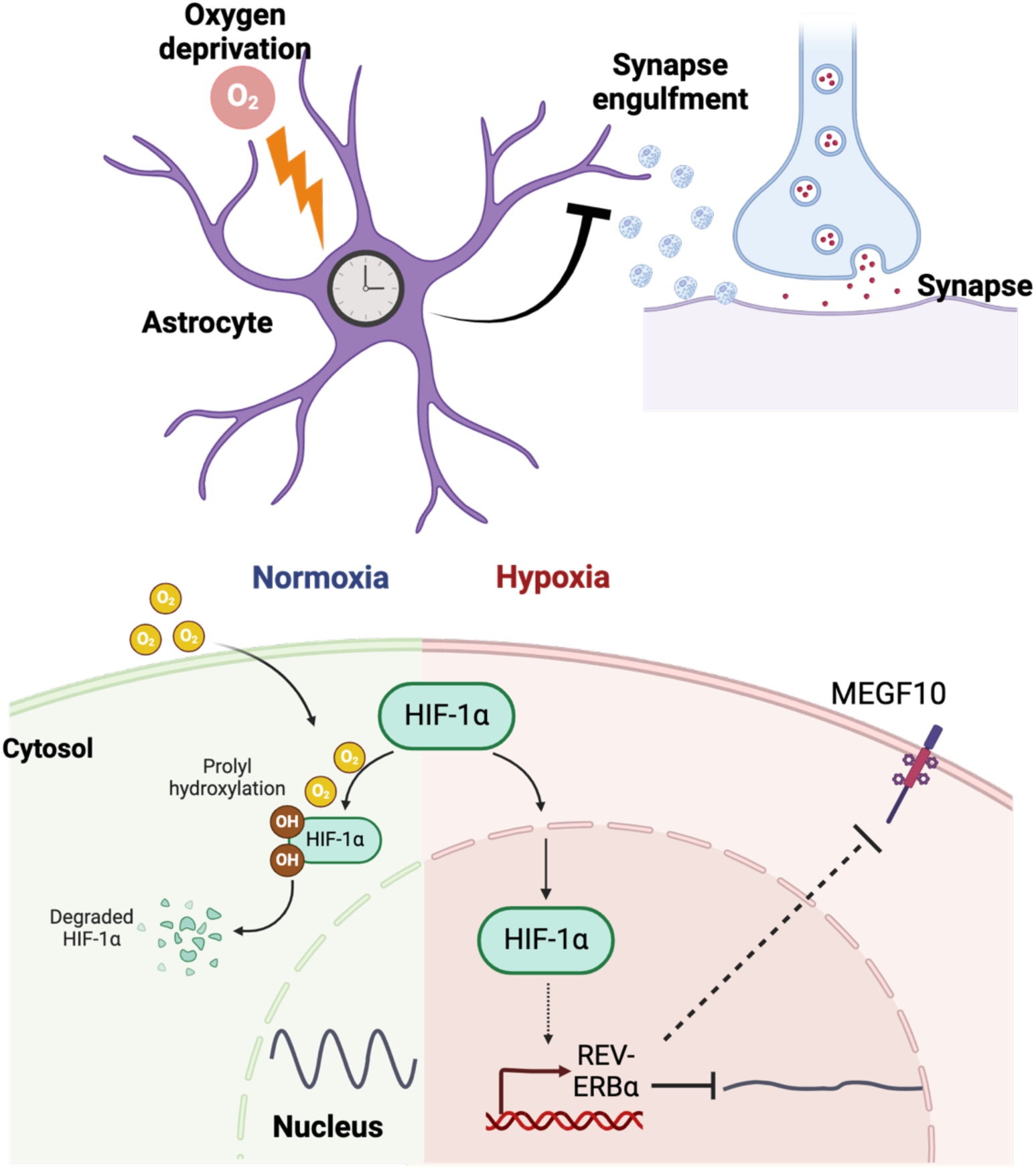
Summary schematic of findings and proposed mechanism of decreased synaptosome engulfment under hypoxia and cellular circadian clock disruptions in astrocytes.

Next, we went on to examine whether pharmacological treatment of astrocytes with SR9011, the enhancer of biological activity of REV-ERBα, induces changes of MEGF10 and MERTK in the absence of hypoxia. We found that indeed, SR9011 treatment also reduces MEGF10 protein level when analyzed by experiment (two-tailed paired *t*-test, P=0.023) and by hiPSC line (two-tailed paired *t*-test, P=0.037) (Figure 5f and g); however, we identified no changes in MERTK protein expression when analyzed by experiment (two-tailed paired *t*-test, P=0.476) or by hiPSC line (two-tailed paired *t*-test, P=0.485) (Figure 5f and h).

The consistent decrease in MEGF10 in both hypoxia- and SR9011-treated astrocytes prompted us to examine whether MEGF10 mediates the synaptosome engulfment defect elicited by enhanced REV-ERBα activity. To address this question, we tested whether overexpression MEGF10 is sufficient to rescue the synapse engulfment defect in astrocytes treated with SR9011. To increase *MEGF10* expression, we first used the CRISPR activation (CRISPRa) system in which cells were transfected with CRISPR-VPR cassettes together with sgRNA pool targeting MEGF10 promoter regions. Cells transfected with CRISPR-VPR cassettes together with sgRNA non-targeting pool (sgRNA scramble) were used as control for comparison. Transfection of CRISPR-VPR cassettes together with sg*MEGF10* significantly increased MEGF10 protein levels in HEK 293T cells (Extended Figure 5a) and mRNA levels in astrocytes compared to sgRNA scramble-transfected cells (Extended Figure 5b).

We next used the CRISPRa system to elevate MEGF10 expression in astrocyte synaptosome engulfment assay. We treated astrocytes with SR9011 or Vehicle only. In SR9011 treated astrocytes, we transfected CRISPR-VPR cassettes along with sgRNA scramble or sgMEGF10 at two doses (2.5nM and 5nM). Consistent with previous experiments, we observed a significant decrease of synaptosome engulfment by astrocytes upon SR9011 treatment compared to Vehicle only treatment when analyzed by experiment (Mixed effect analysis, P=0.008) and by hiPSC line (RM one-way ANOVA, P=0.039) (Figure 5i and j). In the presence of CRISPR activation of *MEGF10* expression, the reduction caused by SR9011 was rescued at both doses of sgRNA, when analyzed by experiment (sgRNA-2.5: Mixed effect analysis, P=0.0003, sgRNA-5: Mixed effect analysis, P<0.0001), and by hiPSC line experiment (sgRNA-2.5: RM one-way ANOVA, P=0.009, sgRNA-5: RM one-way ANOVA, P=0.006) (Figure 5i and j).

Lastly, since MEGF10 expression appears to correlate with REV-ERBα activity, we wondered if MEGF10 expression displays a cyclical expression pattern. To investigate this, we synchronized 10-months hCOs by medium change and collected samples every 6 hr, for a total of 48 hr. We observed that MEGF10 protein levels do oscillate (JTK_Cycle, P=0.007), and, importantly, they do this in an opposite manner of REV-ERBα (JTK_Cycle, P=0.022) (Figure 5k, l and m).

Overall, these results provide strong evidence that synaptosome engulfment defects elicited by elevated REV-ERBα activity are mediated by the decrease of phagocytic receptor MEGF10 expression. In addition, these results demonstrate that MEGF10 expression is negatively regulated by REV-ERBα, a key negative regulator of the circadian clocks.

## DISCUSSION

Neural organoids have become an essential platform for scientific investigation of biological mechanisms underlying neurodevelopmental diseases. In previous work, we demonstrated the power of this platform to model developmental hypoxic cortical gray matter^17,20^, identify cellular phenotypes, define molecular mechanisms of insult and pinpoint pharmacological rescue interventions.

In this manuscript, we studied the effect of hypoxia on hCO-derived human astrocytes at perinatal-equivalent stages on maturation, relevant for hypoxic ischemic encephalopathy. We identified that hypoxia exposure significantly decreases the synaptosome engulfment by astrocytes. Next, we demonstrated the presence of circadian rhythms in control hCOs and their disruption by hypoxia exposure, and showed that this disruption is present in astrocytes but not neurons. We further revealed that one of the mechanisms of synaptosome engulfment defects by hypoxic astrocytes is mediated through decreased expression of MEGF10 phagocytosis receptor, and that this is a direct result of the upregulation of REV-ERBα, a negative regulator of the cellular circadian clock pathway. Finally, we validated the decrease in synapse engulfment by astrocytes with disrupted circadian clocks in vivo, using pharmacological upregulation of REV-ERBα in the hippocampus of an adult mouse model.

This study is the first to demonstrate the presence of circadian rhythms in cortical organoids and to establish a direct mechanistic connection between hypoxia, cellular circadian rhythm disruptions and synaptic engulfment by astrocytes.

The findings in this manuscript are relevant for multiple neurodevelopmental disorders including ASD and ADHD. On one hand, perinatal hypoxic ischemic encephalopathy is a known risk factor for ASD and ADHD^11,14^. Separately, genetic risk factors have identified polymorphisms in circadian clock-related genes (e.g. *CLOCK*, *NR1D1* (*REV-ERBα*), *PER1*, *PER2*) in ASD^32^, and clinical studies consistently confirm the presence of circadian rhythm dysregulations in individuals with ASD^33^.

Based on these data from existing literature and from our current findings, we suggest that the gene-environment interactions, through the presence of genetic predisposition superimposed on hypoxic events during neonatal-early childhood period, significantly contribute to abnormal progression and maturation of circadian clocks in astrocytes, and lead to subsequent altered synaptic pruning and increased risk for ASD and other neuropsychiatric diseases.

We also suggest that the astrocytes are currently a significantly under-recognized contributor to neuronal network sculpting, through significant contributions to synapse pruning, a process historically attributed to microglia. We believe our hypothesis is well supported by the knowledge that the proportion of microglia remains stable during development and maturation (∼ 5% of glia), while astrocytes increase their numbers starting in the second half of pregnancy and eventually become 30-50% of all glia.

Despite all these novel and important findings, we acknowledge our study has several limitations. First, our hCOs are an *in vitro* model and lack the complexity of the developing human brain. However, we did validate the effects of REV-ERBα dysregulation on synaptic engulfment by astrocytes using an *in vivo* adult rodent mouse model within the hippocampus. This suggests that our findings are valid *in vivo* and likely relevant for understanding the role of hypoxic brain injury and/or circadian rhythms disruptions as contributors to the pathophysiology of neurodevelopmental diseases.

Future studies should directly focus on the effects on hypoxia on synapse engulfment *in vivo* and the effects of REV-ERBα expression disruption on synaptic engulfment in neonatal mice brain and more specifically in the cortex. We speculate the findings are going to be replicated in all ages and in all regions of the brain, since this process is so essential and likely evolutionary conserved among ages, brain regions and species. Second, in this study we focus on the acute effects of hypoxia on the astrocytes’ capacity to engulf synaptosomes. Future studies should investigate other functional effects (e.g. cytokine release) and cell-cell interactions, as well as the long-term functional effects following an acute hypoxic event.

Overall, our study is bringing novel mechanistic insights into the effects of hypoxia on essential neurodevelopmental processes (e.g. synapse pruning, circadian disruptions), which have been previously associated with increased risk for neuropsychiatric disorders. The observation of robust circadian rhythms in hCOs is important for the entire scientific community focused on understanding its role in development, injury and senescence.

## METHODS

### Culture of hiPSCs

The hiPSC lines used in this study have been characterized for their pluripotency in previous studies^4,17^ and cultured as previously described. Briefly, hiPSCs were cultured in Essential 8 medium (Life Technologies, A1517001) on vitronectin-(Life Technologies, 14190) coated cell culture dishes. They were passaged every ∼4 days using 0.5 mM EDTA (Life Technologies, 15575) when confluency reaches ∼80%. A total of 4 hiPSC lines derived from 4 individual subjects (2 female and 2 male) were used in this study. Cell integrity was validated using high-density SNP arrays. Mycoplasma was routinely tested using PCR. All hiPSCs and derived hCOs were maintained Mycoplasma free during culture.

### Generation of hCO from hiPSCs

The generation of hCOs from hiPSCs was performed following the protocol described previously^17^. Briefly, hiPSCs were dissociated into single cells using Accutase (Innovate Cell Technologies, AT-104) and seeded into AggreWell plates (STEMCELL Technologies, 34815) at the density of 3 X 10^6^ cells/well in Essential 8 medium (Life Technologies, A1517001) supplemented with 10 μM Rock inhibitor Y-27632 (10 μ M, Selleckchem, S1049). Spheres formed the next day were collected and transferred to ultra-low-attachment 10 cm dishes (Corning, 3262). They were maintained in Essential 6 medium (Life Technologies, A1516401) supplemented with dual SMAD inhibitors SB-431542 (10 μ M, Tocris, 1614), dorsomorphin (2.5 μ M, Sigma-Aldrich, P5499) and Wnt pathway inhibitor XAV-939 (2.5 μ M, Tocris, 3748) for the first 6 days, with every day medium change. From Day 7, hCOs were cultured in Neurobasal A (Life Technologies, 10888) supplemented with B27 (Life Technologies, 12587), 20 ng ml^−1^ epidermal growth factor (R&D Systems, 236-EG) and 20 ng ml^−1^ basic fibroblast growth factor (R&D Systems, 233-FB), with every day medium change for another 8 days, and every other day medium change till Day 25. At this time point, hCOs were switched to neural basal medium supplemented with B27, 20 ng ml^−1^ brain-derived neurotrophic factor (BDNF; Peprotech, 450-02) and 20 ng ml^−1^ NT3 (Peprotech, 450-03) for another 20 days, with every other day medium change. From Day 45 onward, cells were maintained in Neurobasal A medium with B27, with medium change every 3-4 days.

### Enrichment of human astrocytes from hCOs

Ten-months old hCOs (300-350 d) were used for asrtocyte enrichment. hCOs were dissociated as described previously^2,18^. Briefly, ∼20 hCOs were chopped into small pieces using a #10 blade and treated with 30 U/ml Papain (Worthington Biochemical, CAS: 9001-73-4) for 60-90 minutes at 37 °C. Cells were collected and titrated into single cells by pipetting. Astrocytes from dissociated hCO cells were enriched as the HepaCAM^+^ population, using immunopanning for RNA-sequencing or FACS for monolayer culture experiments.

For RNA-sequencing, astrocytes were enriched using immunopanning as previously described^2,19^. Dissociated hCO cells were loaded onto anti-Thy1 (R&D; MAB7335) antibody coated petri dishes (Corning; 351029) to remove neurons. Cells that did not attach to Thy1 dishes were transferred to anti-HepaCAM (R&D, MAB4108) antibody coated petri dishes for astrocytes enrichment. After 15-20 minutes incubation at room temperature, non-attached cells were washed off using DPBS (+Calcium; + Magnesium; Cytiva; SH30264.01), while attached cells were collected following Trypsin-EDTA (Thermo Scientific; 25200114) treatment for RNA isolation.

For synaptosome engulfment assays, astrocytes were enriched using FACS. Dissociated hCO cells were stained with HepaCAM antibody (R&D, MAB4108) at 4 °C for 20 minutes, followed by DPBS (Thermo Fisher Scientific, 14-190-250) washing twice. Secondary antibodies Alexa-Fluor 488 (Thermo Scientific, A-21202) were applied to the cells and incubated at room temperature for 15 minutes, followed by DPBS washing for twice. Cells were filtered through a 40 μm Nylon filter into FACS tubes (CellTreat, 229482) supplemented with live/dead cell dye CytoxBlue (Fisher Scientific, S11348) and proceeded for FACS enrichment. Live cells that are CytoxBlue negative and Alexa Fluor 488 positive cells were gated and collected into neural basal medium supplemented with Rock inhibitor Y-27632 (10 μ M, Selleckchem, S1049). After collection, cells were attached onto poly-D-lysine (Thermo Scientific; A3890401) and Laminin (Thermo Scientific; 23017015) -coated plates at 10, 000 cells/10ul and cultured in neural basal medium.

### Exposure of hCOs and enriched astrocytes to hypoxia

Human cortical organoids (hCOs) or enriched astrocytes derived from 4 hiPSC lines were used for hypoxia exposure. At 10 months (300-350 days) of *in vitro* 3D culture, hCOs or astrocytes enriched at each time points were either maintained in 21% O_2_ (∼ 150 mmHg, 5% CO_2_, 37°C) or were transferred to a C-chamber hypoxia sub-chamber (Biospherix) or HypoxyLab (Oxford) in neural medium for 48 hr. The medium was equilibrated overnight at 30 mmHg, 5% CO_2_ at 37 °C. The level of oxygen in hypoxia chamber (Biospherix) was controlled using a Proox 110 Compact Oxygen Controller and a premixed 5% CO_2_ with balancing N_2_ gas source. The gas levels in HypoxyLab (Oxford) were controlled using built-in gas sensors for O_2_, CO_2_, and N_2_ to achieve stable 30 mmHg and 5% CO_2_ gas levels. After 48 hr, hCOs and enriched astrocytes were immediately collected for analyses.

### Measurement of Oxygen tension in cell culture medium

Cell culture medium for hCOs or astrocytes was exposed to hypoxia using C-chamber hypoxia sub-chamber (Biospherix), HypoxyLab (Oxford), or Spark plate reader (Tecan) following the protocol of hypoxia treatment for the cells as described above. Cell culture medium placed in the control incubator was run in parallel. After 48hr, an optical microsensor of oxygen tension (OXB50, PyroScience) attached to a fiber-optic multi-analyte meter (FireStingO_2_, PyroScience) was immersed into the culture medium to detect oxygen tension, as previously described^17,20^.

### Western blot

Whole-cell protein lysates were prepared from hCO or enriched astrocytes using RIPA lysis buffer (Santa Cruz Biotechnology, sc-24948A) within HypoxyLab after 48 hr of hypoxia exposure. Protein concentration was quantified using Pierce BCA Protein assay kit (Thermos Scientific, 23225). Protein samples were normalized and prepared in 1X LDS Sample Buffer (Invitrogen, 2399420) and 1X Sample Reducing Buffer (Invitrogen, 2353153) with 0.1M DTT. Samples (20 ug/sample/lane) were loaded and run on a NuPage Bis-Tris 4-12% protein gel (Thermo Scientific, 23225), and transferred onto a 0.45 um PVDF membrane (Fisher Scientific, IPFL00010). Membranes were blocked using 5% Milk in 1X PBST for 1 hr at room temperature (RT) and incubated with primary antibodies against β-ACTIN (Cell Signal Technology, Anti-Mouse, Clone 13E5, 4970S), HIF1-α (Cell Signaling Technology, Anti-Rabbit, Clone D2U3T, 14179S), REV-ERBα (Abnova, Anti-Mouse, Clone 4F6, H00009572-M02) for overnight at 4C. The next day, membranes were washed and then incubated with HRP-linked secondary antibodies: anti-Rabbit (Cell Signaling Technology, 7074P2) or anti-Mouse (Cell Signaling Technology, 7076P2) for 1 hr at RT. Protein bands were developed using Pierce ECL Western Blotting Substrate (Thermo Scientific, 32209) or SuperSignal West Femto Maximum Sensitivity Substrate (Thermo Scientific, 34095) and imaged using ChemiDocX Imager. Quantification was performed using ImageStudioLite with normalization to β-actin controls.

### Viral labeling of dissociated hCO and enriched astrocytes

Dissociated hCOs or enriched astrocytes were plated in 24-well culture plates (Corning, 353047) and incubated with lentivirus overnight at 37°C with 5% CO_2_. The next day, fresh neural medium was added to replace the virus. hGFAP::GFP infected cells were cultured for another 3-5 days for imaging analysis. PER2:LUCIFERASE infected cells were cultured for another 7-10 days for luminescence recording.

### Immunocytochemistry

Monolayer cells cultured in 8-well chamber slides (Thermo Scientific, 154941) were fixed in 4% PFA (Electron Microscopy Sciences, 15710) for 15 min at 4C, washed with PBS twice and blocked in Blocking Buffer containing 10% donkey serum (NDS, Millipore-Sigma, S30-M) with 0.3% Triton X-100 (Millipore Sigma, T9284-100ML), for 1 hr at RT. Then cells were incubated with primary antibodies at appropriate dilutions (listed in Extended Table 6) in Blocking Buffer overnight at 4C. The next day, secondary antibodies were applied for staining at 1:1000 dilution in PBS for 1 hr at RT, followed with mounting in Aqua-Poly/Mount (Polysciences, 18606-5).

### Astrocyte synaptosome phagocytosis assay and quantification

Synaptosome purification and *in vitro* phagocytosis assays were performed as previously described ^2,7^. Briefly, synaptosomes were purified by Percoll (Cytiva, 17-0891-02) gradient from adult Sprague-Dawley rat brains (Charles River) and incubated with pHrodo Red (Life technology, P3660). The hCO-astrocytes were seeded in separate culture plates for control or hypoxia treatment and labeled with hGFAP::GFP lentivirus as above, and maintained for 3-5 days. Enriched astrocytes were FACS-sorted using HepaCAM antibody and seeded in different 8-well Chamber Slides for control and hypoxia exposure. CellTracker CMFDA (Thermo Scientific, C2925) was supplemented to culture medium right before imaging to label enriched astrocytes. Then one plate was continually maintained in the incubator with normal oxygen level, and the other plate was transferred to a hypoxia chamber for a total of 48 hr, followed by imaging. After the first 24 hr, 12.5 μg purified synaptosomes were supplemented into culture medium, and images were collected using the same parameters across conditions per experiment using Confocal Microscope (Zeiss LSM980). 5-15 fields, 10X magnification, were set up using multi-position imaging per well to cover all the cells.

To quantify synaptosome engulfment, images were first encoded using ReNamer and then analyzed blindly by a different person. Images were imported to Imaris and converted to imz format. pHrodo-Red signal and GFP signal were measured using the Volume function in Imaris. Each channel was thresholded separately for individual images using consistent criteria based on intensity across images. Overall pHrodo-Red volume was normalized to GFP volume per image. After Imaris quantification, image names were decoded for comparisons between conditions and statistical analysis.

### RNA sequencing and analysis

RNA from immunopanned astrocytes or neurons from 10-month hCOs was isolated using RNAeasy Micro Kit (Qiagen, 74004). Bulk RNA sequencing libraries were prepared using the NEBNext Ultra II kit with PolyA Selection and sequenced using the Illumina HiSeq 4000 System to a minimum of 30 million paired-end reads per sample. Fastq files were first trimmed using Trimmomatic and mapped to the hg38 human genome reference using STAR aligner with the paired end option selected. We used FeatureCounts to assemble transcripts and generate raw count matrices. We generated alignment quality-control metrics with RSeQC v4.0.0. Following count matrix generation, we used DESeq2 to normalize matrices and to determine differential gene expression statistics. We identified a total of 1,480 differentially expressed genes in 10-months astrocytes, based on two cutoffs: Log_2_ Fold Change ≥ 0.5 and FDR < 0.01. We used a Gene Ontology analysis online tool (http://geneontology.org/) to identify GO biological processes from DEGs. PANTHER Overrepresentation Test (Released 20221013) was performed with Fisher’s Exact Test and False Discovery Rate (FDR) was calculated. Database used at the time of analysis was GO Ontology database DOI: 10.5281/zenodo.6399963 Released 2022-03-22. For maturation comparisons, we adopted the gene lists enriched in progenitors and mature astrocytes from primary human astrocytes identified from previous studies^15^. TPM values of genes from 4 iPSC lines at two differentiation stages (6-month and 10-month) were pulled out from our RNA-seq results and presented as heatmap in Extended Figure 1b.

### PER2:LUCIFERASE assay in hCOs and rhythmicity analysis

Intact hCOs at 10-months were transduced overnight with PER2:LUCIFERASE lentivirus (gift from Erik Herzog Lab). One week after viral transduction, we transferred 3 hCOs to each well of a white 96-well plate with transparent bottom (Nunc, ThermoFisher 165306) and performed media changes on samples for synchronization. 0.5 μM Luciferin was supplemented in medium and luminescence was recorded using Tecan SPARK with environmental control at 37°, 5% CO_2_ and 21% O_2_ for control condition, or 5% CO_2_ and 3.5% O_2_ (P*O_2_* equivalent to ∼30mmHg) for hypoxia condition. Luminescence measurements (counts/second) were recorded at the gap of 10 min for a total 48 hr. Relative luminescence was calculated by normalizing to the 1st luminescence value from 2 hr after media change (Timepoint 13, T13). To analyze rhythmicity of recordings, normalized luminescence values at one hour gap (T13, T19, T25…to T289) were uploaded to NiteCap for JTK_Cycle analysis. The dataset from each well containing 3 hCOs were analyzed as individual repeats.

### RNAscope assay and quantification

Astrocytes were FACS sorted as described above and plated in 8-well Lab-Tek II®CC2 Slide (Thermo Scientific, 1054941). Within 2-3 days after plating, astrocytes were exposed to control or hypoxia for 48 hr as described previously, then fixed with 4% PFA for 15 mins followed by processing for RNAscope assay as described^20^. Briefly, slides were dehydrated using 50%, 70%, 100%, 100% ethanol for 5 min each. Then hydrogen peroxide was applied for 10 min. Primary antibody of GFAP (Dako, Z033429) was applied to the slides and incubated at 4°C overnight. The next day, slides were fixed in 4% PFA for 30 min followed with Protease III treatment for another 10 min. Slides were baked for 45 min and proceeded with RNAScope Multiplex Fluorescent V2 Co-detection kit (ACD Biosciences) following manufacturer’s protocol. The probes used were: *REV-ERBα* (808441-C2), *CLOCK* (853011-C2). Imaging was performed using Leica SP8 confocal. Maximum projections were generated using LASX software and further analyzed using ImageJ (Fiji) following instructions from the kit manufacturer (ACD Biosciences).

Astrocytes area was selected as R.O.I by GFAP staining in each image. 25-100 cells were selected per image depending on cell density on the image. The number of dots representing probed transcripts were quantified using ImageJ and exported for statistical analysis.

### siRNA knockdown of *REV-ERBα* and synaptosome engulfment assay

siRNA scrambled control and two siRNAs targeting different sequences in Exon 1 of *REV-ERBα* were designed and synthesized by Dharmacon (si*REV-ERBα*-1, J-003411-09-0002) and Santacruz (si*REV-ERBα*-2, sc-61458). To check the efficiency of *REV-ERBα* knockdown, 25nM siRNAs were transfected into HEK 293T cells using 0.6μL RNAimax reagent (Thermo Fisher Scientific, 13778075) with 50 μl Opti-MEM (Thermo Fisher Scientific, 31985062). Reduction of *REV-ERBα* in HEK 293T cells treated with both siRNAs targeting *REV-ERBα* compared to scrambled control was confirmed using qRT-PCR. To knockdown *REV-ERBα* in astrocytes, enriched astrocytes using HepaCam FACS were plated in 8-well chamber slides as described above. After 2-3 days recovery, siRNAs transfection was performed in astrocytes using the same condition as in HEK 293T cells. Two days after transfection, medium change was performed, and chamber slides were transferred to HypoxyLab for hypoxia treatment at 30mmHg following synaptosome engulfment assay using enriched astrocytes and Confocal imaging as described above.

### Pharmacological treatment

Astrocytes were seeded and labeled with hGFAP::GFP lentivirus as above. SR9011 (Cayman Chemical Company, 11930) was supplemented to the cells at final concentration 10 μM, along with enriched synaptosomes 24 hr before imaging for both normal and hypoxic conditions. Medium with SR9011 was replaced with neural medium before imaging. Synaptosome phagocytosis was imaged and quantified as described above.

### Mice

All experiments with mice were conducted according to articles approved by the Institutional Animal Care and Use Committee (IACUC) from Korea Advanced Institute of Science and Technology (KAIST), and we followed ethical regulations. Mice were maintained in a standard plastic cage (29 cm (L) x 17.5 cm (W) x 18 cm (H)). Wild-type C57BL/6 mice were purchased from Samtaco and Deahan BioLink.

### In vivo astrocyte phagocytosis assays

At 8-12 weeks, pAAV::hSyn-Synaptophysin1-mCherry-EGFP (Expre) construct was packaged into Adeno-associated virus serotype 9 (AAV9) and 300 μl of virus was injected into mouse brain CA3 (AP: -1.9, ML: +-2.36, DV from pia mater: -2.20, virus titer = 5*10^12 GC/ml) as previously reported^15^. SR10067 (Tocris Bioscience), and vehicles were formulated with 10% dimethyl Sulfoxide (DMSO, LPS Solution) and 90% Corn oil (Sigma-Aldrich, Cat. No. C8267). For western blot, qPCR, and astrocyte phagocytosis experiments, mice were reared in light/dark (L/D) cycle with 12 hours each at least for a week followed in dark/dark (D/D) cycle from a day before SR10067 injection. SR10067 was intraperitoneally injected with 30 mg/kg dosage for three subsequent days at CT6, at which *NR1D1*, the target of SR10067, expression level is highest^14,16^. Mice were sacrificed at CT0 after three times injection of SR10067.

Mice were anesthetized with 1.2% avertin (20 μl/g) by intraperitoneal injection. Anesthetized mice were perfused with Dulbecco’s Phosphate Buffered Saline (DPBS, Welgene), followed by 4% paraformaldehyde (PFA, Wako Chemicals). The prepped brain was maintained in 4 % PFA overnight in 4 °C followed by 30% sucrose in DPBS at least 24 hours for dehydrating. The dehydrated brain was embedded in OCT compound (Leica) and sectioned with 30 μm by Cryostat microtomes (Leica).

For immunostaining, the brain sections were embedded in blocking solution (4% Bovine serum albumin in 0.3% Triton X-100 in DPBS) in a room temperature (RT) for an hour. After blocking, brain slices were stained with primary antibody, a rabbit anti-S100β (Abcam 1:1000) and a rat anti-LAMP2 (Abcam, 1:500), in blocking solution overnight in 4 °C followed by 5 times wash with PBST (0.1% Tween20 in DPBS). The washed slices were stained with secondary antibody, with Alexa fluorophore 647-conjugated donkey anti-rabbit antibody (Abcam, 1:250), and Alexa fluorophore 405-conjugated donkey anti-rat antibody (Abcam, 1:250) in blocking solution for 2 hours in RT, followed by 5 times wash with PBST. The stained slices were mounted onto adhesive slide glass (Matsunami) and treated with Trueblack (Biotium) to quench autofluorescence from lipofuscin. The samples were then covered with mounting media (VECTASHIELD) for confocal microscopy imaging.

Confocal images were taken from the CA1 SR region with LSM 880 microscopy (Zeiss, 40x oil immersion optical lens). Five stacks of images (1 μm per stack) were acquired, and two images were taken from one mouse. Images were split by EGFP (488), mCherry (594), and S100β (647) channels for processing in Image J (or FIJI). Among split images, mCherry (594) channel was subtracted by EGFP (488) channel to acquire mCherry-only puncta. The subtracted image and S100β (647) channel were conducted by colocalization analysis by DiAna plugin in ImageJ. The area of the colocalized mCherry-only puncta with S100 β were measured, and was normalized by the area of S100β. To prevent mCherry-alone puncta from a distortion by a virus expression level, the measured area was further normalized by EGFP area.

### Overexpression of *MEGF10* using CRISPR-VPR activation

All components of single guide RNA (sgRNA) were designed and synthesized by Dharmacon, including pooled 4 CRISPR RNAs (crRNAs) targeting promoter region of *MEGF10* (P-014897-01-0005), crRNA non-targeting control (U-009500-10-05) and trans-activating CRISPR RNA (tracrRNA) (U-002005-05). *GFAP*::dCas9-VPR (CRISPRa) plasmid was designed by VectorBuilder. To examine the efficiency of MEGF10 overexpression by CRISPRa system, HEK 293T cells were co-transfected with CRISPRa plasmid (200ng) and pooled MEGF10 crRNAs using 7.5 μl Lipofectamine 3000 (Thermo Fisher Scientific, L3000001) following the manufacturer’s instructions. Increased MEGF10 expression in the presence of MEGF10 crRNAs compared to crRNA non-targeting control were confirmed using qRT-PCR or Western blot. Primer sequences for qPCR and sequences of MEGF10 crRNAs and qPCR primers are provided in Extended Tables 7 and 8, respectively. To increase MEGF10 expression in astrocytes, MEGF10 crRNAs or non-targeting controls with dCas9-VPR plasmid (200ng) were transfected into HepaCam enriched astrocytes seeded in 8-well chamber slides using the lipofectamine LTX reagent (0.6μl) and plus reagent (0.2μl, Thermo Fisher Scientific, A1262) containing 40μl Opti-MEM following the manufacturer’s protocol. Two doses of crRNAs, 2.5nM and 5nM were used based on minimal toxicity to the cells and efficiency of MEGF10 overexpression. Two days after transfection, fresh culture medium change was performed and proceeded with validation of MEGF10 expression or synaptosome engulfment assay.

### STATISTICAL ANALYSES

Data are presented as mean ± S.E.M. unless otherwise indicated. Distribution of the raw data was tested for normality; statistical analyses were performed as indicated in figure legends. Sample sizes were estimated empirically or based on power calculations. Blinding was used for analyses comparing control and treatment samples.

## Supporting information

Extended Table 1 - lines used for each experiment

DEGs in 10-month astrocytes associated with hypoxia exposure

Gene ontology in 10-month astrocytes associated with hypoxia exposure

Log2fold changes and padj values of genes in each GO category

DEGs in 10month neurons associated with hypoxia exposure

Primary antibody list

Primers for RT-qPCR

Sequences for CRISPRa synthetic crRNAs

## ACKNOWLEDGMENTS

We thank the members of the NeoPasca lab and other collaborators for constructive feedback and or editing the manuscript (alphabetical order: Tudor Braicu, Yusuke Hori, Madison House, Emily Gurwitz, Geana Marte, Yuki Miura, Sergiu P. Pasca, Mayuri Vijay Thete, Deniz Yagmur Urey, Andrea Navarrette Vargas, Sejin Yoon, Erik Herzog). This project was supported by Stanford Maternal & Child Health Research Institute (MCHRI) Postdoctoral Fellowship (to L. L.) and Bill and Melinda Gates Foundation. Figures were created with BioRender.com.

## DATA AVAILABILITY

Gene expression data are available in the Gene Expression Omnibus (GEO) under accession number GSE255588. All data and other codes are available from the corresponding author upon request.

## AUTHOR CONTRIBUTIONS

L.L. performed experiments, data analysis and manuscript writing. J.C. and S. H., A.P., N.R., and D.N. contributed to experimental design, data collection and analyses. C.S. and W.C. performed in vivo mouse experiments, data analysis and manuscript editing. S.M. and S.L. performed data analysis for the synaptosome engulfment experiments. A.V. and S.S. contributed to data analyses for RNA-sequencing, figure generation and manuscript writing. J.L. and J.K. contributed to RNAscope experiments and imaging. D.R. and E.G. contributed to circadian rhythm experiments analysis. A.M.P. supervised the work and wrote the manuscript with input from all authors.

## COMPETING INTEREST STATEMENT

Stanford University was granted a patent that covers the generation of region-specific brain organoids. A.M.P is listed as inventor for U.S Patent number: 10,494,602 B2. \

---

**EXTENDED FIGURE 1.**
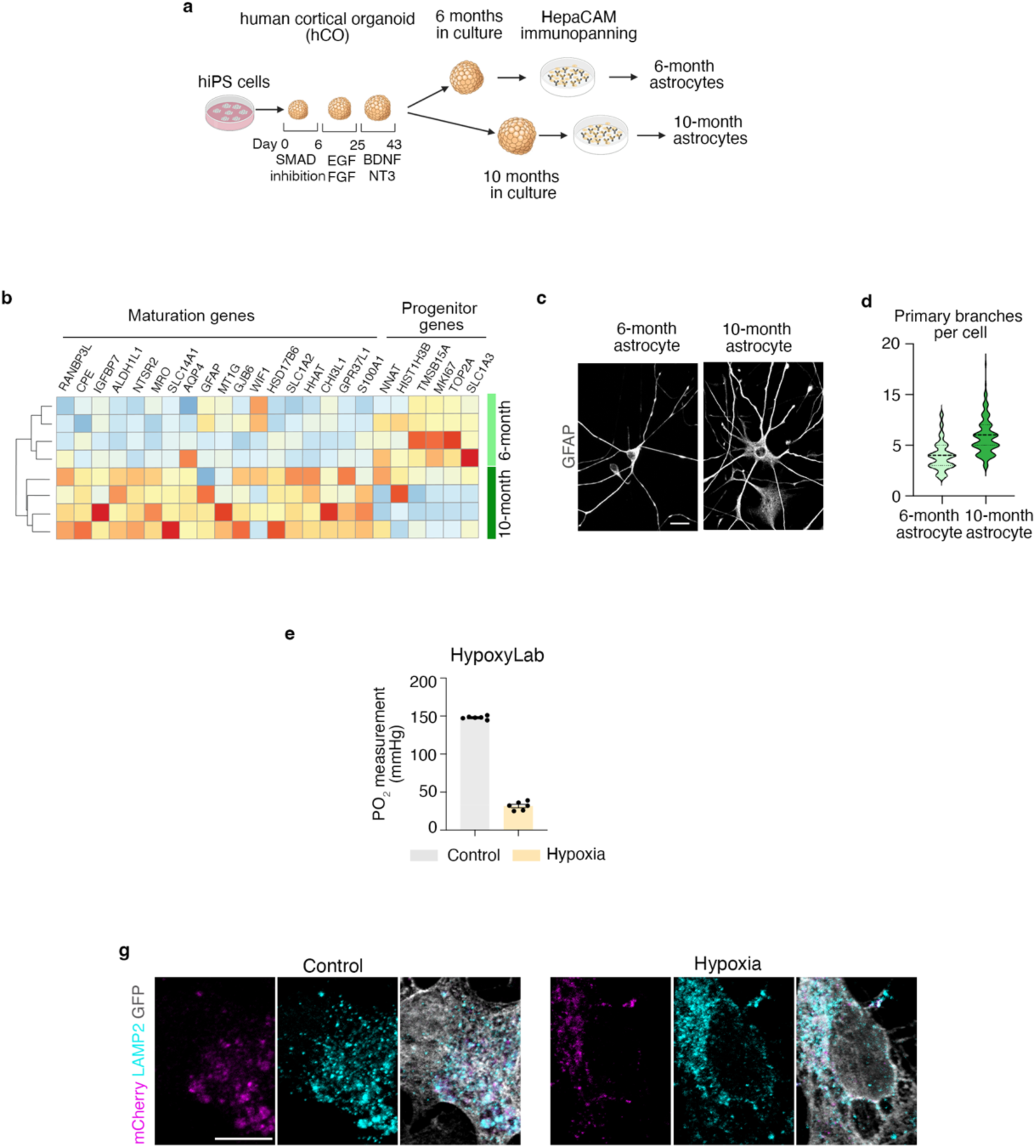
Parameters of astrocyte maturation and technical validation of assays. **(a)** Schematic of experimental design for purification of astrocytes from hCOs at 6 months and 10 months, through immunopanning using HepaCAM antibody; **(b)** Heatmaps of TPM values of genes associated with astrocyte maturation between 6- and 10-months hCO-derived astrocytes. Each row represents one hiPSC line; **(c)** Immunostaining with GFAP showing morphological differences between 6 and 10 months astrocytes; scale bar 20 μm; **(d)** Quantification of primary branches in 6-months and 10-months astrocytes; **(e)** Measurement of oxygen tension in medium of cells in control and hypoxia conditions (n=6 independent measurements); **(f)** Co-localization of synaptosomes engulfed into astrocytes with lysosome marker LAMP2 in control- and hypoxia-exposed astrocyte; scale bar: 10 μm.

**EXTENDED FIGURE 3.**
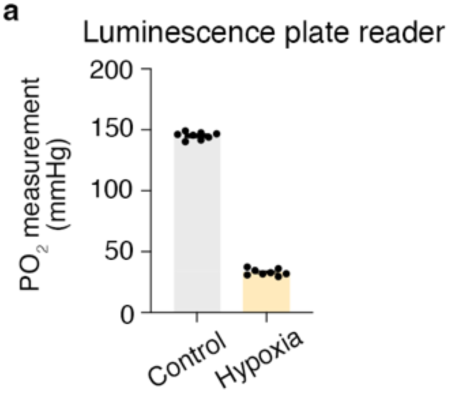
Oxygen tension measurements in the luminescence plate reader environment. **(a)** Measurement of oxygen tension in the medium of cells in control and hypoxia conditions (n=8 independent measurements) in the Tecan Spark plate reader used for luminescence recording.

**EXTENDED FIGURE 4.**
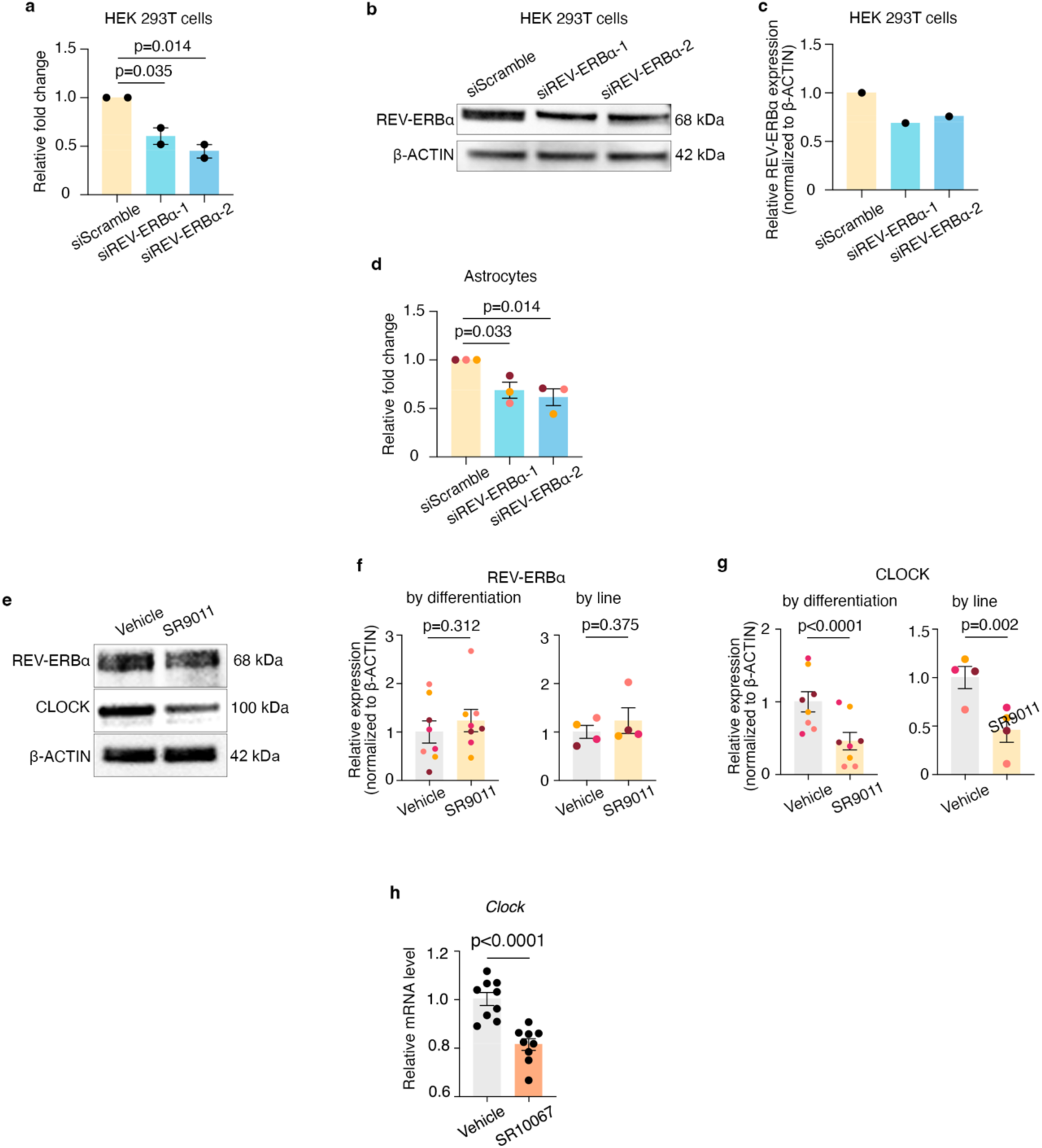
Effects of REV-ERBα activity level modification on astrocytes. **(a)** Quantification of *REV-ERBα* gene expression changes following addition of siScramble, si*REV-ERBα*-1 and si*REV-ERBα*-2 in HEK 293T cells (one-way ANOVA; si*REV-ERBα*-1: P=0.035; si*REV-ERBα*-2: P=0.014); **(b)** Representative Western blot images of REV-ERBα and β-ACTIN proteins in the presence of si*REV-ERBα*-1 and si*REV-ERBα*-2 in HEK 293T cells; **(c)** Quantification of REV-ERBα protein level relative to β-ACTIN in the presence of siScramble versus si*REV-ERBα*-1 and si*REV-ERBα*-2 in HEK 293T cells; **(d)** Quantification of *REV-ERBα* gene expression changes in siScramble versus si*REV-ERBα*-1 and si*REV-ERBα*-2 conditions in 10-months hCOs-derived astrocytes from 3 hiPSC lines (siScramble vs si*REV-ERBα*-1: one-way ANOVA, P=0.014; siScramble vs si*REV-ERBα*-2: one-way ANOVA, P=0.033); **(e)** Representative Western blot images of REV-ERBα, CLOCK and β-ACTIN proteins in vehicle versus SR9011-treated 10-months hCOs; **(f)** Quantification of REV-ERBα protein levels relative to β-ACTIN in vehicle versus SR9011-treated 10-months hCOs analyzed by experiment (paired Wilcoxon test; P=0.312) and by hiPSC line (paired Wilcoxon test; P=0.375); **(g)** Quantification of CLOCK protein levels relative to β-ACTIN in vehicle versus SR9011-treated 10-months hCOs, analyzed by experiment (two-tailed paired *t*-test; P<0.0001) and by hiPSC line (two-tailed paired *t*-test; P=0.002); **(h)** Quantification of Clock gene expression level in the mouse brain upon vehicle versus SR10067 treatment (two-tailed unpaired *t*-test; P<0.0001). Each color of dots represents one cell line for in vitro experiments and one mouse brain sample for in vivo Clock qPCR; data are presented as mean ± s.e.m.

**EXTENDED FIGURE 5.**
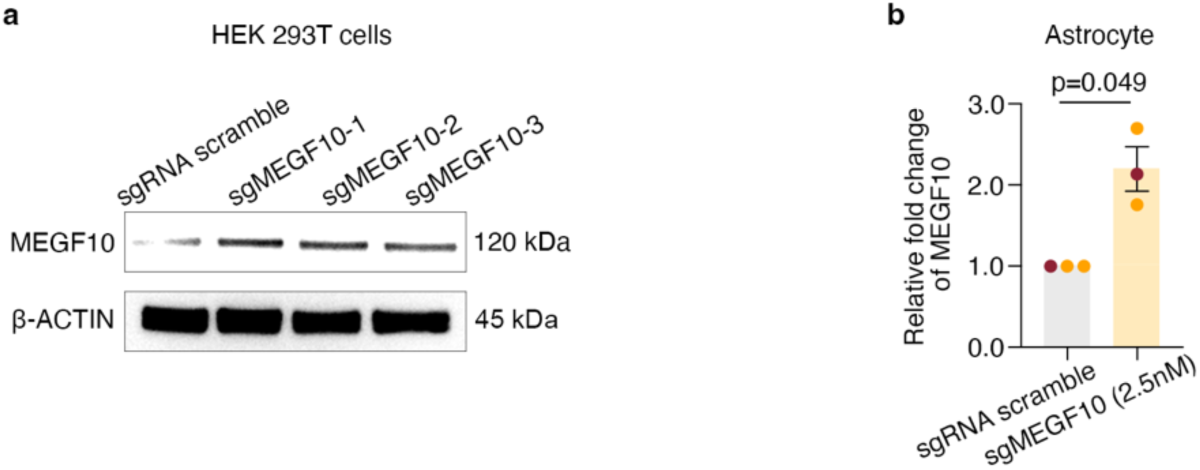
**(a)** Representative Western blot images of MEGF10 and β-ACTIN proteins in the presence of sgRNA non-targeting pool (sgRNA scramble) and sgRNA pool targeting MEGF10 promoter (sgMEGF10) in HEK 293T cells. sgMEGF10-1, -2, and -3 are three independent transfections in HEK 293T cells; **(b)** qRT-PCR quantification of MEGF10 mRNA levels in 10-months hCO-derived astrocytes in the presence of sgRNA scramble, or sgMEGF10 transfected at 2.5nM.

## Notes

### Competing Interest Statement

The authors have declared no competing interest.

### Summary of Updates

list of authors was changed to reflect corresponding author as last listed author

